# Time Dependent Dihedral Angle Oscillations of the Spike Protein of SARS-CoV-2 Reveal Favored Frequencies of Dihedral Angle Rotations

**DOI:** 10.1101/2023.01.02.522466

**Authors:** Oscar Bastidas

## Abstract

The spike protein of SARS-CoV-2 is critical to viral infection of host cells which ultimately results in COVID-19. In this study we analyze the behavior of dihedral (phi and psi) angles of the spike protein over time from molecular dynamics and identify that the oscillations of these dihedral angles are dominated by a few discrete, relatively low frequencies in the 23-63 MHz range with 42.96875 MHz being the most prevalent frequency sampled by the oscillations. We further observe that upon tallying the populations of each individual frequency for all residues along the frequency spectrum, there is a regular alternation between high and low population counts along the increasing frequency values in the spectrum. This alternation of the counts becomes less pronounced and ultimately stabilizes as the frequency values increase. These observations thus suggest a regularity and propensity in the spike protein’s dihedral angles to avoid similar oscillation population counts between vicinal frequencies. We also observe that for amino acids that are least abundant in the S protein, there are certain frequencies at which the dihedral angles never oscillate, in contrast to relatively abundant amino acids that ultimately cover the entire spectrum. This suggests that the frequency components of dihedral angle oscillations may also be a function of position in the primary structure: the more positions an amino acid is found in, the more frequencies it can sample. Lastly, certain residues identified in the literature as constituting the inside of a druggable pocket of the spike protein, as well as other residues identified as allosteric sites, are observed in our data to have distinctive time domain profiles. This motivates us to propose residues from our dynamic data, with similar time domain profiles, which may be of potential interest to the vaccine and drug design communities, for further investigation. Thus our findings indicate that there is a particular frequency domain profile for the spike protein, hidden within the time domain data, and this information, perhaps with the suggested residues, might provide additional insight into therapeutic development strategies for COVID-19 and beyond.

## Introduction

As of December 2022, the World Health Organization reports that SARS-CoV-2, the pathogen responsible for COVID-19, has killed over 6.6 million people worldwide since the start of the pandemic [1]. This deadly virus infects the epithelial cells lining the respiratory tract by fusing its spike (S) protein with the Angiotensin Converting Enzyme 2 (ACE2) receptor on the host cell [2–4]. Although vaccines have been instrumental in curbing death rates, their efficacy has been observed to wane after 5-8 months and this, along with the rise of troubling new variants, creates a concern over the possibility of continuing infections of COVID-19 [5]. Given the S protein’s critical role in the initial phases of infection, it is one of the most important drug targets and so understanding its architecture and mechanics remains a relevant research topic. The S protein is thus composed of three identical chains where each chain is made up of two subunits (S1 and S2) [6]. While the S2 subunit forms the fusion peptide, the S1 subunit is in turn composed of four domains: the N-terminal domain (NTD), C-terminal domain 1 (CTD1), C-terminal domain 2 (CTD2) and the receptor binding domain (RBD) [7, 8]. Binding between the S protein and ACE2 thus begins with a conformational change of the RBD domain from the “down” position to the “up” position. Previous studies investigating various aspects of the S protein’s motions and mechanics include work that identified a latch region on the S protein that appears to regulate these “down” to “up” conformations [9]. Another study has identified centripetal motion of the RBD domain upon binding with ACE2 along with specific residues that played an important role in those motions [8]. Other work has taken the approach of considering the effects of dihedral angles associated with “down” to “up” conformational changes with another study obtaining insight into the entropy of residues most amenable to binding, by similarly looking at dihedral angles [10, 11]. Although these approaches have yielded very important information in the fight against COVID-19, a dedicated characterization of the dihedral angles themselves is still lacking that specifically probes phenomena pertinent to time-informed dihedral angle fluctuations, such as, frequencies of oscillations and identifying potential dominant frequencies throughout the S protein. Consequently, the precedent currently remains modest for the questions posed by our work which motivates our present efforts.

For this work, our approach consists of analyzing dihedral angle data (phi and psi), as a function of time, obtained from molecular dynamics simulations from our previous study of the trimeric S protein (PDB ID: 6VSB) [9]. The RBD domain of chain A for this structure is in the “up” position. For every residue, we analyze frequency data obtained from the time domain to look for any frequencies that have a dominant/majority presence in individual chains, among amino acids (i.e. among all ARGs), as well as the whole protein. We also curate this data to see if there are any frequencies that uniquely persist for any of the twenty amino acids throughout the simulation period. Lastly, we attempt to identify correlations between each residue’s motility in Euclidean space and its dihedral angle fluctuation variations. Phi and psi angle data are thus analyzed separately.

## Materials and Methods

### Molecular Dynamics

The trajectory of protein motions was obtained from our previous work [9] which used explicit solvent molecular dynamics simulations of the SARS-CoV-2 S protein using the NAMD2 program to carry out the simulation for a total of 200 nsec after equilibration with trajectories being written every nanosecond. Using CHARMM-Gui [12], the protein was thus explicitly solvated with TIP3P water molecules and the CHARMM36m force field was selected. Missing residues in the experimental structure were added and disulfide bonds and glycosylated sites were also included. The simulation was carried out using the NPT ensemble which maintains the number of simulated particles, pressure and temperature constant. We used the Langevin piston method to maintain a constant pressure of 1 atm and we employed periodic boundary conditions for a water box simulation volume along with the particle mesh Ewald (PME) method with a 20 Å cutoff distance between the simulated protein and the water box edge. The integration time step was 2 femtoseconds and our simulation was conducted under physiological conditions (37 °C, pH of 7.4, physiological ionic strength with NaCl ions, LYS and ARG were protonated, HIS was not).

### Signal Processing

Due to there being no sudden forces applied to the protein during the simulation that could result in steady state changes to the vibrational frequencies, for every residue, the Fourier transform was obtained from the phi vs. time and psi vs. time dihedral angle data (coming from the molecular dynamics simulations), using the fast Fourier transform algorithm from the JDSP library for the Java programming language. Positive absolute values of frequencies were thus calculated for the Fourier transform and the sampling frequency was once per nanosecond. Frequency domain data consequently spanned from 0 MHz to 496.09375 MHz, just under the Nyquist/folding frequency of 500 MHz set by our fixed sampling rate (Nyquist frequency being half the sampling rate).

### Data Analysis

In order to identify frequencies that might be particularly abundant in the S protein, each time a peak appeared at a given frequency in the Fourier transform for an individual residue, it was tallied (the location of a peak at a given frequency in the individual residue’s spectrum being indicative that the dihedral angle possessed that frequency as a component of its oscillations). Peaks were identified as local maxima in the frequency domain plots and so the DC component frequency at 0 MHz (typically a large value relative to the spectral peaks in our data) was disregarded. Such a tally was carried out as three independent analyses: 1) at the amino acid level, 2) at the individual chain level and 3) for the whole protein. Tally information was then plotted to visualize the abundance of the frequencies for each of the three cases. Next, root mean square fluctuations (RMSF) were calculated for every residue, and in order to quantify the tendency of dihedral angles to step outside of their respective mean (potentially a foreshadow to protein structural changes), population standard deviations were calculated for both phi and psi for each individual residue over the simulation period. These two metrics were then plotted (RMSF vs. phi/psi standard deviation) to see if there were any correlations observed between the two quantities. The standard deviations for phi and psi were then separately plotted as histograms to infer the spread of this variation. Lastly, in order to identify if any frequencies were unique to an amino acid (i.e. unique to all LYS in the protein), we carried out SQL inner table joins of the frequency data where peaks were specifically present, organized by amino acid (the table joins being responsible for identifying which frequency values were present across all the data specific to the amino acid being analyzed i.e. across all LYS). The complete data set is found in the Supplemental Information material which is available upon request.

## Results and Discussion

Tallying all of the frequencies across the spectrum first of all revealed that the most abundant frequency, throughout the entire protein, was 42.96875 MHz, for both phi and psi dihedral angles, and the frequencies of 62.50000 MHz and 27.34375 MHz were observed as the second and third, respectively, most prevalent frequencies likewise for both phi and psi. Secondly, the profile of the plotted tally counts also showed that there was a regular alternation of high counts followed by low counts along the frequency spectrum and this difference in counts gradually became less pronounced as the frequency increased. This profile behavior was also observed for the plotted tally counts for just the individual chains, and to a lesser degree, for the amino acid level as well, within a given chain (see Figure 1). The third observation was that for amino acids that had a relatively low abundance in the protein (HIS, MET and TRP), at the amino acid level within any given chain, it was observed that there were frequencies at which the dihedral angles (both phi and psi) never oscillated (see Figure 2). There were between 9 and 15 of each of these three low abundance amino acids per chain, where each chain had over 1000 residues. These un-accessed frequencies consequently appear as gaps in those plotted tallies. At the level of the whole protein (i.e. when tallies from all chains were combined), however, HIS in particular was observed to have populations of frequency counts that covered the whole spectrum for both phi and psi (Figure 3). MET and TRP on the other hand still had un-accessed frequencies/gaps at the whole protein level for the phi dihedral angle, which were found at the lowest frequencies, but were able to cover the entire spectrum with psi (Figure 3).

**Figure 1:**
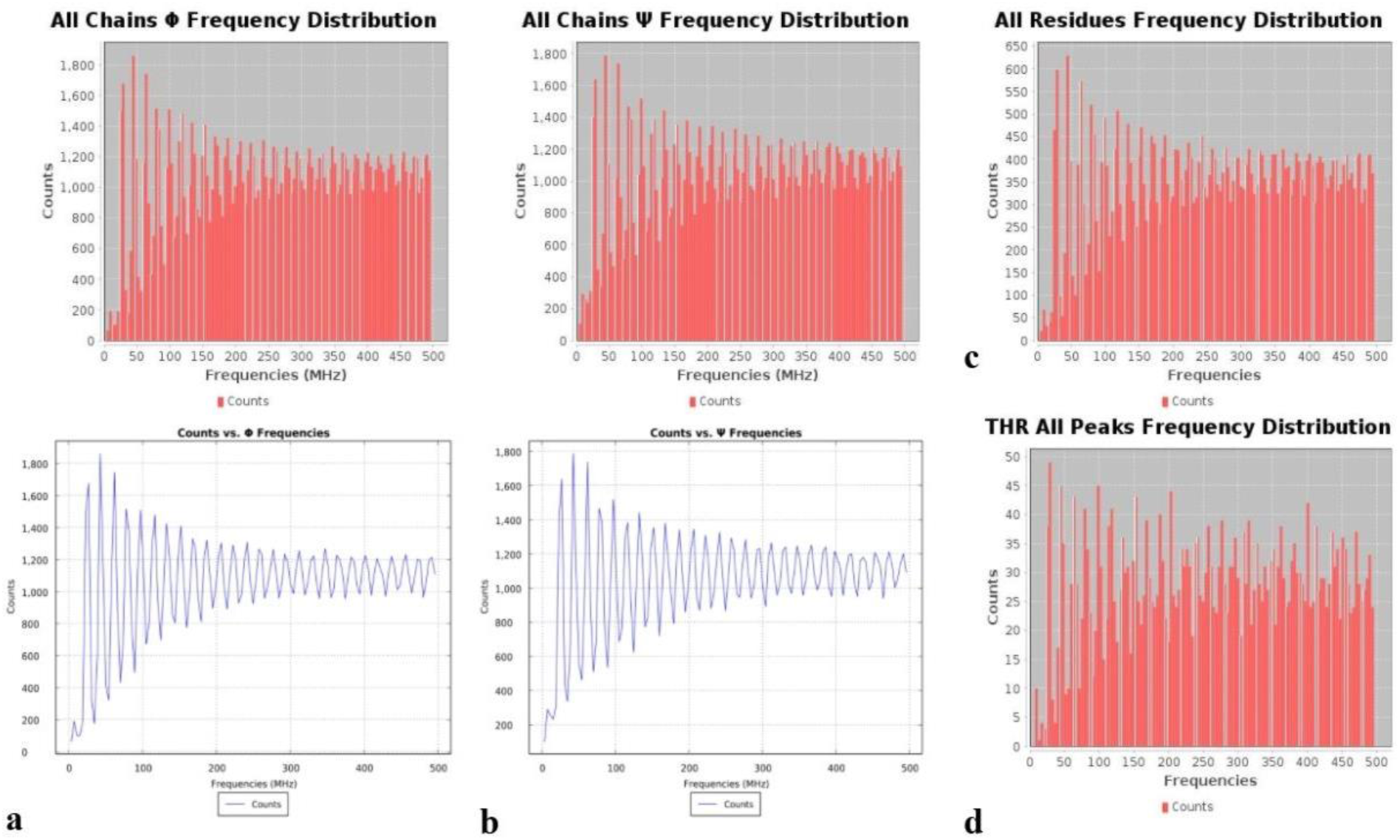
Tallied frequencies for all residues across all three chains for: a) the phi dihedral angles showing bar chart in top panel and line plot in bottom panel (for ease in seeing profile), b) the psi dihedral angle showing bar chart in top panel and line plot in bottom panel (for ease in seeing profile). c) Tallied phi frequencies for all residues in the A chain and d) tallied phi frequencies for all threonines in the A chain (as an example of tally profile at the amino acid level).

**Figure 2:**
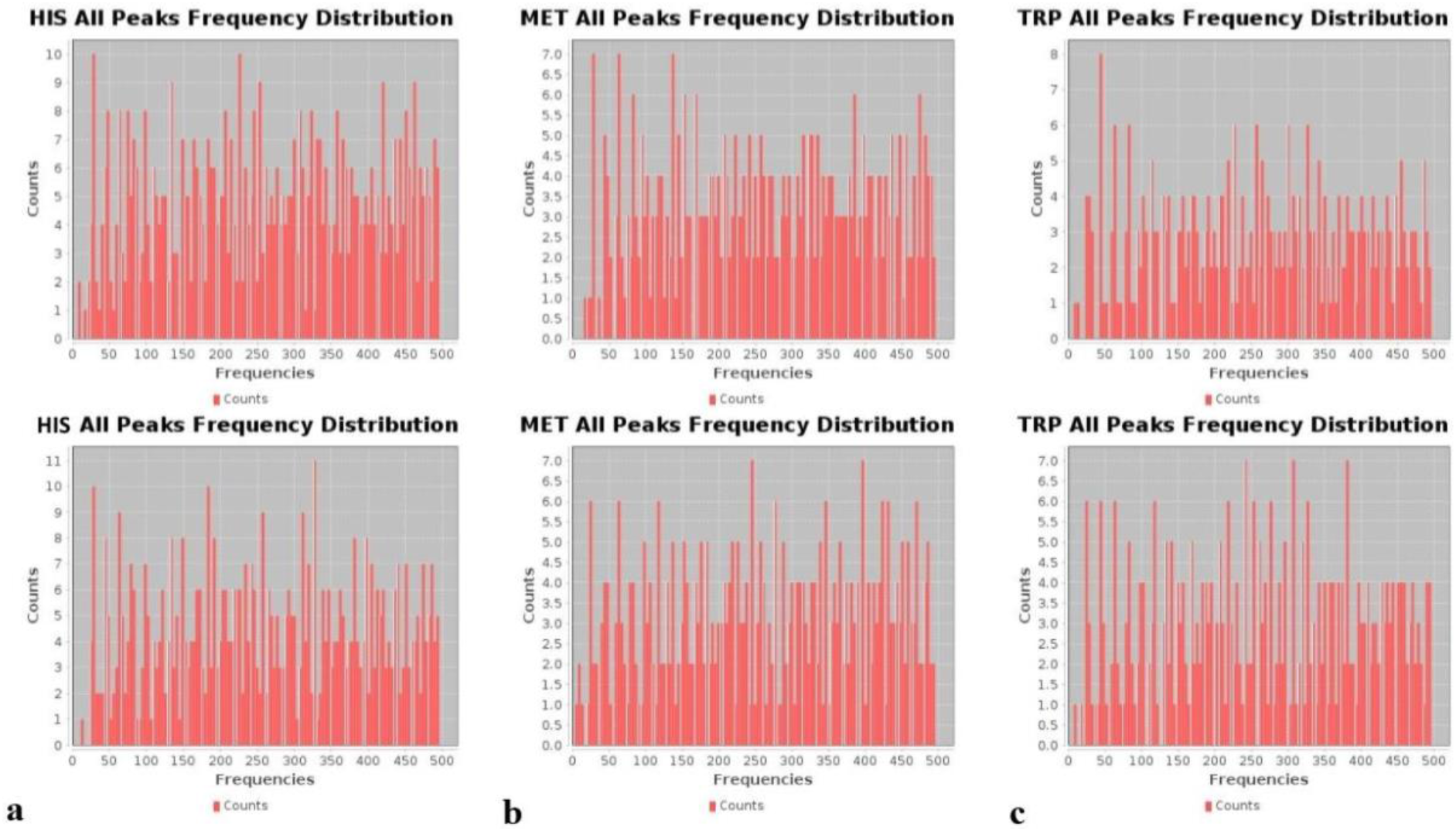
Frequency tallies showing un-accessed frequencies for: a) all histidines in the A chain, phi and psi in the top and bottom panels respectively, b) all methionines in the B chain, phi and psi in the top and bottom panels respectively, c) all tryptophans in the C chain, phi and psi in the top and bottom panels respectively.

**Figure 3:**
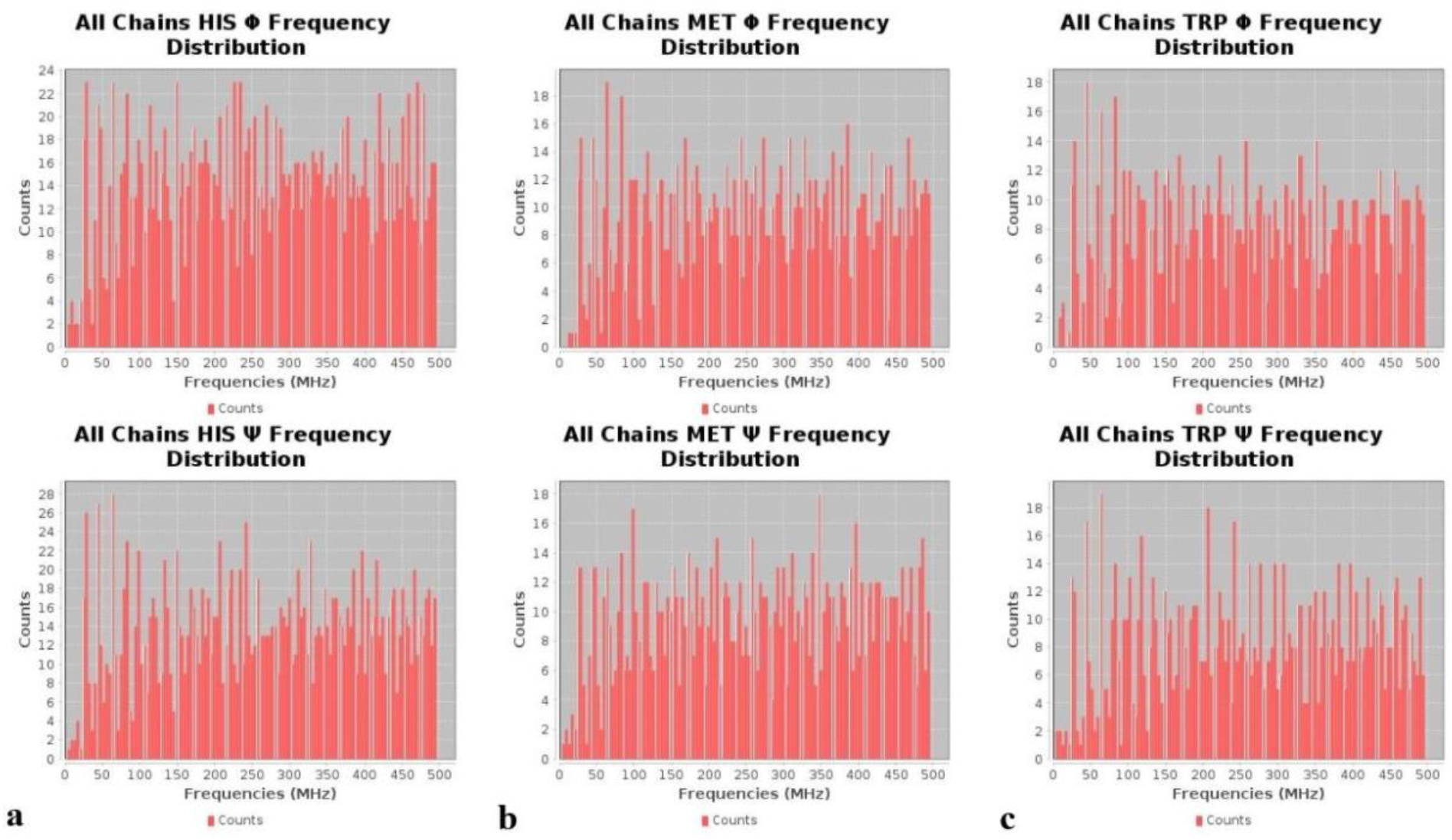
Tallied frequencies across all three chains for: a) all histidines, phi and psi tallies in the top and bottom panels respectively, b) all methionines, phi and psi tallies in the top and bottom panels respectively, c) all tryptophans, phi and psi tallies in the top and bottom panels respectively

For each individual chain, plotting all of the residue’s RMSFs as a function of the standard deviations of their psi dihedral angles revealed that there was a larger quantity of residues with relatively large RMSF values corresponding to the larger values of the psi standard deviations (over 90°) compared to phi (Figure 4). This was in contrast to the results observed for the phi standard deviations where there were fewer residues at these higher values of the phi standard deviations and what residues were present, they had relatively lower RMSF values (Figure 4). The identity of the residues for both of these cases for standard deviations greater than 90° is thus shown in Table 1, specifically, for those residues with the top twenty RMSF values (phi, however, consistently did not have a full set of 20 amino acids at this upper range of standard deviations). Histogram inspection of the distribution of the values of the standard deviations, for both phi and psi, also revealed that both dihedral angles had a positive skew profile in their values (Figure 5). Looking at each amino acid throughout each individual chain (i.e. all LYS in chain A, then in chain B, then chain C separately) revealed that there were no frequencies whatsoever that persisted throughout the simulation for a given amino acid.

**Figure 4:**
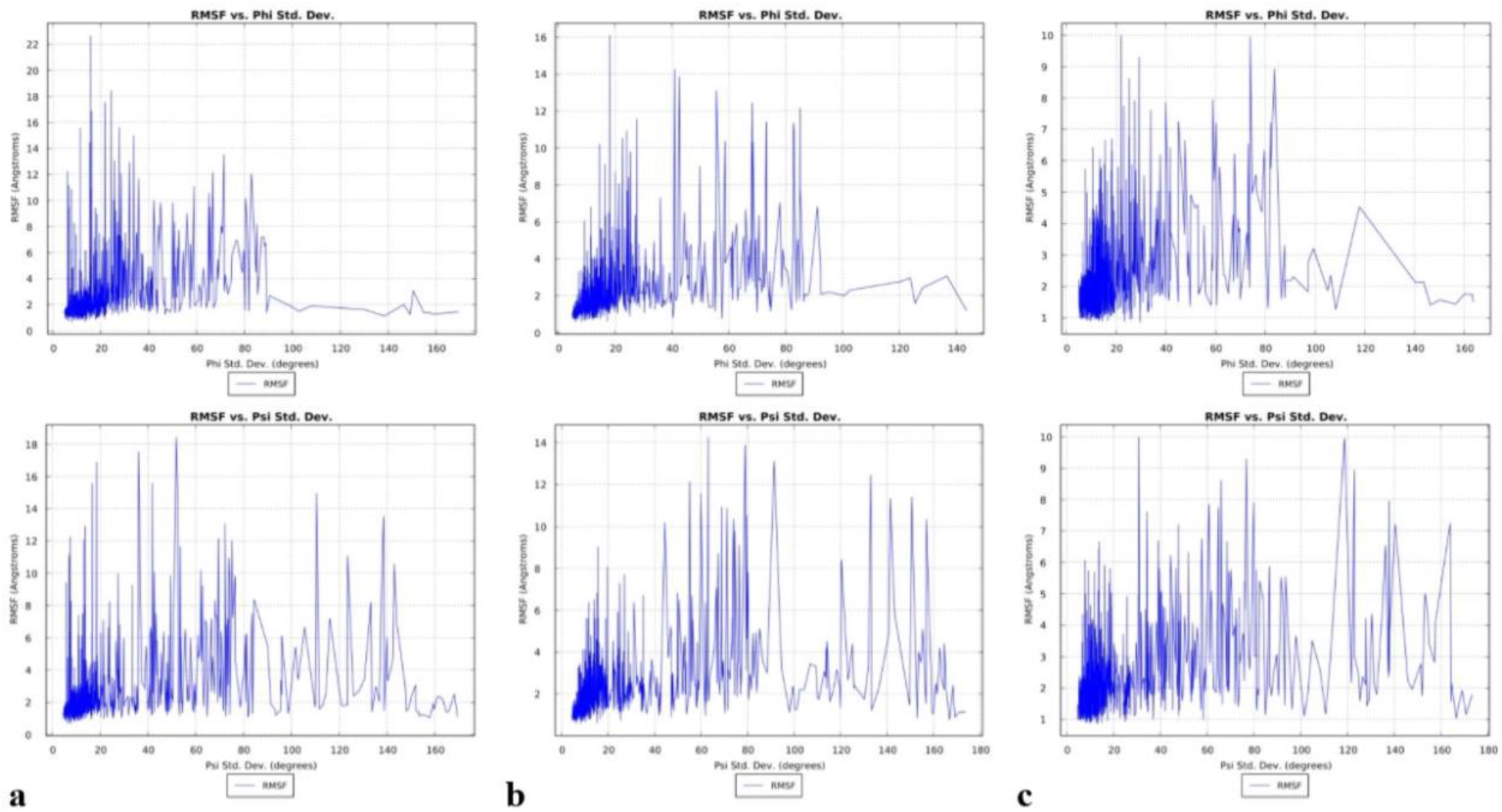
RMSF vs. dihedral angle standard deviation for: a) chain A, phi and psi in the top and bottom panel respectively, b) chain B, phi and psi in the top and bottom panel respectively and c) chain C, phi and psi in the top and bottom panel respectively

**Table 1:** Identity of residues with the top 20 RMSF values for dihedral angle standard deviations over 90°for: a) chain A phi, b) chain A psi, c) chain B phi, d) chain B psi, e) chain C phi and f) chain C psi. Phi angles did not have as many residues above a standard deviation of 90° compared to psi.

| a |  |  |  | b |  |  |  | c |  |  |  | d |  |  |  | e |  |  |  | f |  |  |  |
| --- | --- | --- | --- | --- | --- | --- | --- | --- | --- | --- | --- | --- | --- | --- | --- | --- | --- | --- | --- | --- | --- | --- | --- |
| Amino Acid | Amino Acid # | RMSF (Å) | Phi Std. Dev. | Amino Acid | Amino Acid # | RMSF (Å) | Psi Std. Dev. | Amino Acid | Amino Acid # | RMSF (Å) | Phi Std. Dev. | Amino Acid | Amino Acid # | RMSF (Å) | Psi Std. Dev. | Amino Acid | Amino Acid # | RMSF (Å) | Phi Std. Dev. | Amino Acid | Amino Acid # | RMSF (Å) | Psi Std. Dev. |
| GLY | 891 | 3.073 | 150.726 | SER | 477 | 14.976 | 110.670 | SER | 256 | 6.825 | 91.248 | SER | 683 | 13.111 | 91.494 | GLY | 1099 | 4.535 | 117.911 | GLY | 181 | 9.945 | 118.758 |
| THR | 500 | 2.684 | 90.701 | GLY | 252 | 13.527 | 138.673 | SER | 98 | 3.537 | 92.306 | ASP | 839 | 12.450 | 133.075 | THR | 941 | 3.208 | 99.492 | LYS | 150 | 8.934 | 122.996 |
| GLY | 89 | 2.010 | 146.756 | GLY | 482 | 11.073 | 123.635 | GLY | 550 | 3.076 | 136.751 | GLY | 842 | 11.405 | 150.773 | GLY | 381 | 2.786 | 97.382 | SER | 254 | 7.951 | 137.810 |
| PHE | 1109 | 1.944 | 99.038 | GLY | 682 | 10.586 | 143.153 | SER | 591 | 2.972 | 123.962 | GLY | 682 | 11.342 | 141.485 | GLU | 324 | 2.358 | 106.367 | GLY | 447 | 7.250 | 163.994 |
| SER | 746 | 1.914 | 108.012 | GLY | 75 | 8.209 | 133.294 | GLY | 1099 | 2.765 | 120.257 | GLY | 838 | 10.328 | 156.917 | ASN | 1125 | 2.313 | 91.400 | GLY | 446 | 7.211 | 140.348 |
| GLY | 593 | 1.627 | 130.430 | GLY | 485 | 8.066 | 137.636 | GLY | 548 | 2.422 | 128.144 | GLY | 257 | 8.405 | 120.450 | GLN | 836 | 2.206 | 90.607 | GLY | 75 | 6.543 | 136.219 |
| VAL | 227 | 1.505 | 103.000 | GLY | 257 | 7.493 | 138.830 | GLY | 107 | 2.295 | 102.508 | ALA | 27 | 5.835 | 143.333 | GLY | 103 | 2.147 | 143.795 | GLY | 184 | 6.339 | 141.429 |
| GLY | 550 | 1.457 | 169.528 | LYS | 150 | 7.194 | 116.148 | SER | 605 | 2.205 | 95.462 | GLY | 181 | 5.090 | 155.467 | ALA | 520 | 2.138 | 140.490 | HSD | 625 | 5.555 | 93.595 |
| GLY | 548 | 1.402 | 157.169 | GLY | 446 | 6.956 | 144.142 | GLY | 103 | 2.100 | 92.363 | GLY | 184 | 4.842 | 140.915 | GLY | 891 | 1.863 | 105.180 | ALA | 626 | 5.522 | 91.693 |
| GLY | 103 | 1.388 | 155.179 | GLY | 447 | 6.659 | 105.651 | ASN | 450 | 2.007 | 100.810 | GLY | 72 | 4.507 | 114.191 | THR | 109 | 1.837 | 97.259 | GLY | 72 | 5.002 | 153.155 |
| GLY | 107 | 1.284 | 158.869 | ALA | 942 | 6.120 | 96.030 | GLY | 593 | 1.600 | 125.560 | GLY | 832 | 4.391 | 164.787 | GLY | 89 | 1.764 | 160.160 | ALA | 27 | 4.458 | 154.234 |
| LEU | 546 | 1.245 | 149.395 | GLY | 72 | 5.998 | 140.044 | GLY | 89 | 1.205 | 143.696 | GLY | 476 | 4.388 | 125.216 | GLY | 593 | 1.751 | 163.270 | ALA | 263 | 4.446 | 134.978 |
| GLY | 1059 | 1.141 | 138.717 | ASP | 442 | 5.474 | 90.189 |  |  |  |  | GLY | 252 | 4.188 | 162.856 | GLY | 548 | 1.571 | 150.302 | PHE | 464 | 4.418 | 92.688 |
|  |  |  |  | THR | 73 | 5.425 | 101.810 |  |  |  |  | GLY | 447 | 4.176 | 113.987 | GLY | 550 | 1.506 | 163.777 | SER | 256 | 4.372 | 130.514 |
|  |  |  |  | ASN | 137 | 4.520 | 142.442 |  |  |  |  | GLY | 446 | 3.987 | 153.751 | GLY | 107 | 1.445 | 156.637 | GLY | 838 | 4.224 | 127.916 |
|  |  |  |  | GLY | 261 | 3.452 | 130.674 |  |  |  |  | GLY | 75 | 3.539 | 154.669 | GLY | 1035 | 1.416 | 146.405 | GLY | 257 | 4.091 | 157.588 |
|  |  |  |  | THR | 333 | 3.406 | 102.933 |  |  |  |  | ASN | 164 | 3.445 | 107.007 | PHE | 1109 | 1.280 | 108.362 | GLY | 842 | 3.931 | 134.412 |
|  |  |  |  | PHE | 1121 | 3.368 | 95.727 |  |  |  |  | GLY | 1124 | 3.335 | 132.111 |  |  |  |  | ALA | 475 | 3.739 | 137.992 |
|  |  |  |  | GLY | 496 | 3.350 | 140.442 |  |  |  |  | GLY | 261 | 3.303 | 109.569 |  |  |  |  | ALA | 262 | 3.664 | 98.027 |
|  |  |  |  | GLY | 891 | 3.073 | 152.079 |  |  |  |  | ARG | 634 | 3.256 | 94.674 |  |  |  |  | LYS | 444 | 3.515 | 104.967 |

**Figure 5:**
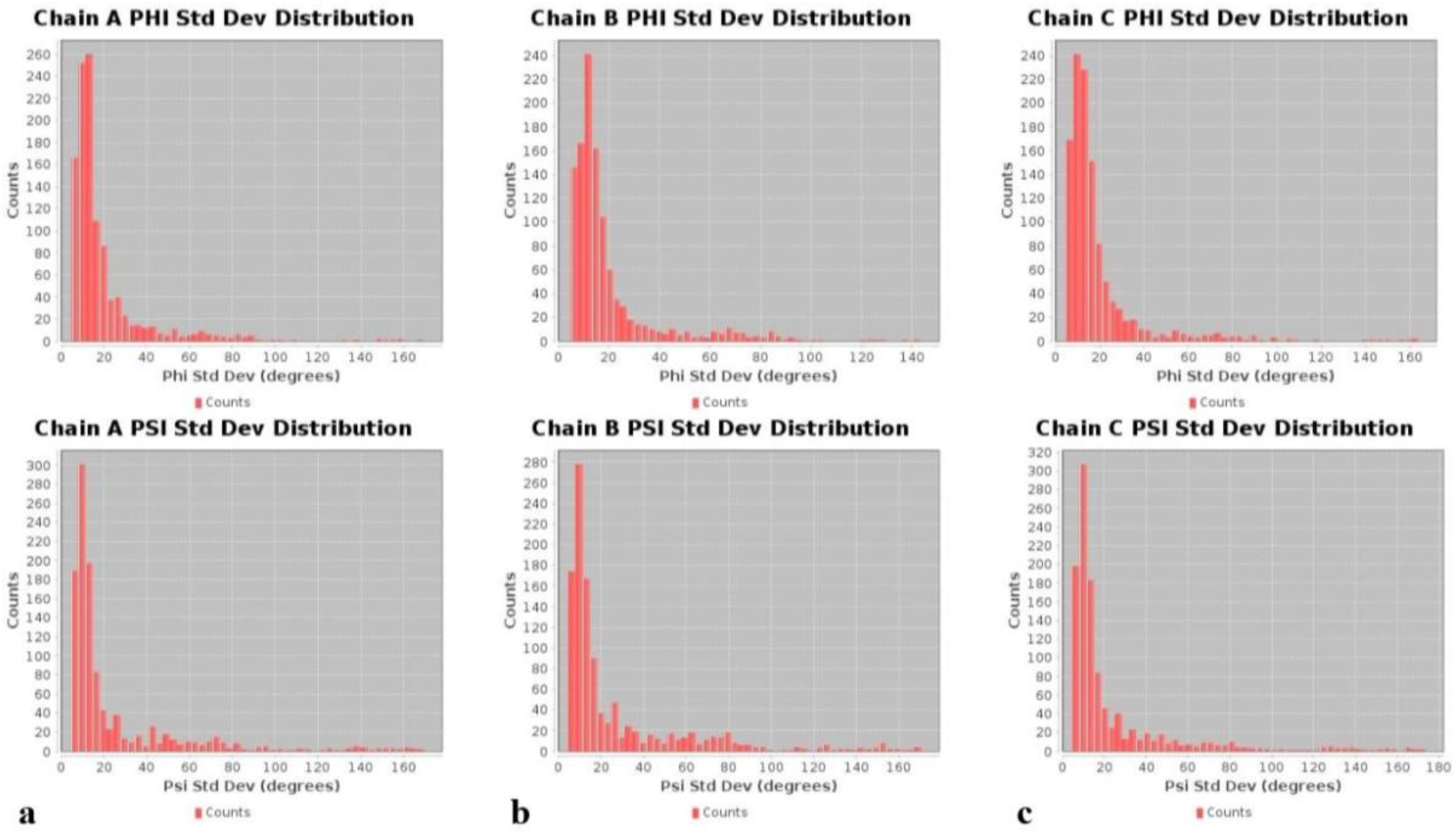
Dihedral angle standard deviation distributions showing positive skew profile for: a) chain A, phi and psi in the top and bottom panel respectively, b) chain B, phi and psi in the top and bottom panel respectively and c) chain C, phi and psi in the top and bottom panel respectively. All histogram distributions done with 50 bins.

Several groups have previously identified residues important to either drug binding, allosteric effects, or both, for the S protein. According to work by Alvarado et al., LYS369 and PHE377 appeared to have important allosteric properties since binding of luteolin to these residues in the S protein induced an intense allosteric effect [13]. Verkhivker et al. reported that GLU406, ASN439, LYS417 and ASN501 were centers of allosteric interactions implicated in mediating long-range communications in the binding of the S protein with ACE2 [14]. Of these residues, LYS417, ASN501 along with the additional residue of GLU484, were further identified to be important interacting centers that provide mutants at these positions with better binding affinity to ACE2. Xue et al. additionally identified PHE329 and PHE515 as residues critical to centripetal motions of the RBD domain upon binding with ACE2 [8]. Drew and Janes also identified a druggable pocket and thus reported the amino acids lining this pocket [15]. Our own data showed that the dihedral angle vs. time profiles (phi, psi or sometimes both) for these very residues had a distinctive profile that consisted of either a dihedral angle rapidly assuming a value far from the baseline, then immediately returning back to baseline, or the baseline would change very suddenly (Figure 6). We note there were no distinctive frequency domain profiles for these residues, however. These unique time domain profiles were in contrast to those observed for most other residues whose dihedral angle values instead oscillated within a single fixed bandwidth of dihedral angle values (Figure 6). Consequently, we present a list of some other residues which were identified in our dynamic data as having similar profiles, for further investigation as potential residues of interest to the drug design community (Figure 7 & Table 2). Solvent accessible surface area (SASA) data for each residue is also provided to give preliminary insight into each residue’s potential accessibility to a drug-like compound.

**Figure 6:**
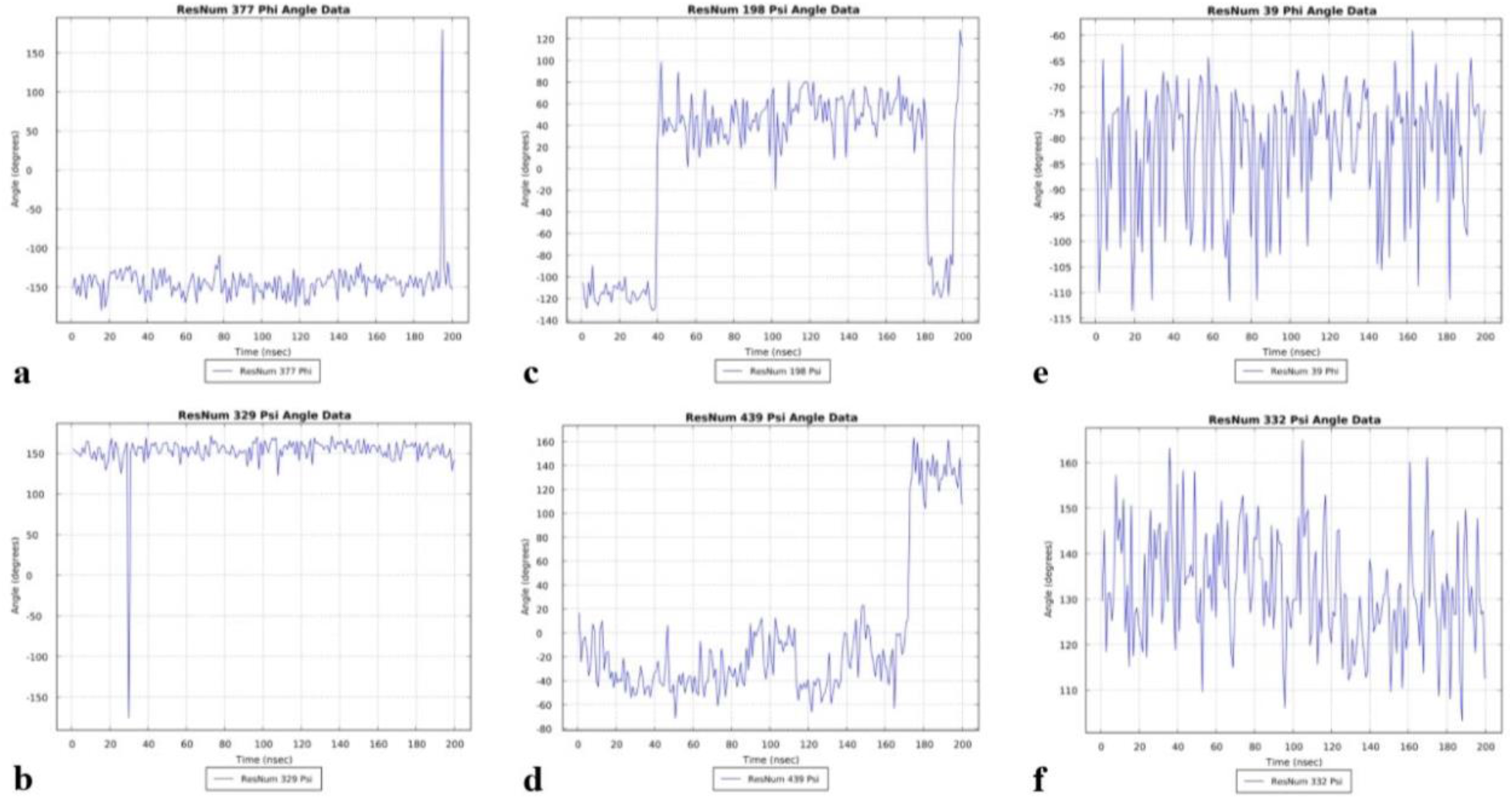
Dihedral angle values vs. time for residues identified in literature as either important to allostery or the formation of a druggable pocket: a) PHE 377 chain C phi [13], b) PHE 329 chain B psi [8], c) ASP 198 chain C psi [15], d) ASN 439 chain A psi [14]. Dihedral angle values vs. time for most other residues: e) PRO 39 chain A phi, f) ILE 332 chain B psi.

**Figure 7:**
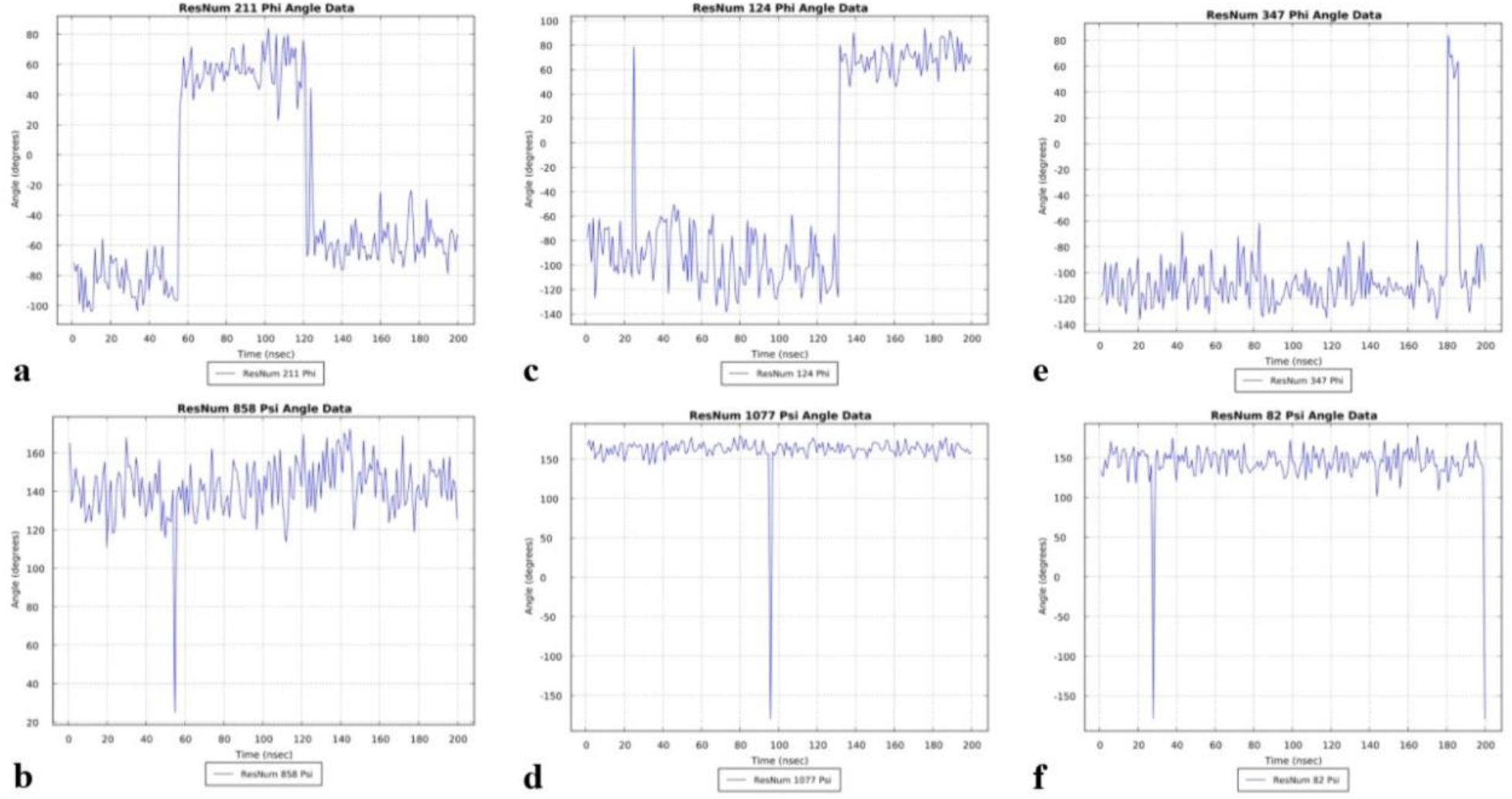
Dihedral angle vs. time data for additional residues from our data set with time domain profiles similar to those residues previously identified as relevant to allostery and drug pocket formation: a) ASN 211 chain A phi, b) LEU 858 chain A psi, c) THR 124 chain B phi, d) THR 1077 chain B psi, e) PHI 347 chain C phi, f) PRO 82 chain C psi

**Table 2:**
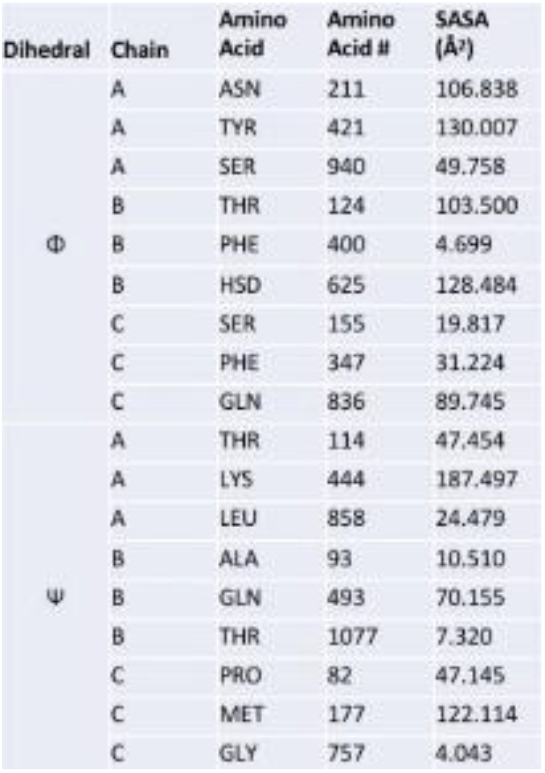
Additional residues with time domain profiles potentially indicative of some relevance to additional allostery or drug pocket formation. SASA data reported in Angstroms squared.

## Conclusions

From the tally of the frequency populations, the most favored frequency components for phi and psi dihedral angle oscillations in the S protein appeared at relatively low frequencies in the 23-63 MHz range with 42.96875 MHz being the most sampled frequency component for the entire protein and individual chains. The profile of the frequency counts also appears to indicate that dihedral angles sample their oscillation frequencies so that they avoid comparable quantities of population counts between frequencies close to each other, resulting in the high-low alternation of the tallies seen along the frequency spectrum (Figure 1). Given that the difference between population counts is relatively large at lower frequencies, but the difference gradually decreases as the frequency increases, the most sampled frequencies are consequently found at the lower end of the spectrum. The observations that the least abundant amino acids in the protein (HIS, MET and TRP) never sampled certain frequencies throughout the simulation, also suggests that the specific frequency components sampled by dihedral angle oscillations might be a function of the position of the amino acid in the protein. We hypothesize this because amino acids that were, in contrast, more abundant (and hence were more distributed throughout the protein’s sequence), had dihedral angle frequency population counts that spanned the entire spectrum and left no gaps at various frequencies (Figure 2). This consequently appears to suggest that the more positions an amino acid occupies in a protein’s primary structure, the more frequencies it would be able to sample in each individual residue’s oscillations. Furthermore, because the un-accessed frequencies were not identical across the three chains for any of these three amino acids (despite the three chains being identical in primary structure), we suggest that the component frequencies sampled by a residue’s dihedral angles might also be a function of the micro environment created by other non-bonded residues within some radial distance to the amino acid in question. Though the chains are identical, their spatial orientation relative to each other would be different since the RBD domain on chain A is in the “up” position, and the other two chains’ RBD domains are in the “down” position (and our simulation was of the isolated protein so no binding partners were present to observe domain position changes). Our data also suggests that residues that have a tendency to experience sudden and wide variations of their psi dihedral angle values outside their mean (i.e. residues that have high psi standard deviations), also seem to be more motile in Euclidean space as indicated by the general positive correlation of those residues’ high psi standard deviation values with their relatively high respective RMSF values (data for high phi standard deviations suggests the opposite behavior). Lastly, residues identified by previous groups as being important to allosteric function and/or drug binding had distinctive time domain profiles in our data. Although our study focused on the S protein isolated (in contrast to many of the aforementioned studies where it was bound to ACE2), we propose the possibility that the time domain profiles we observed might be a preview into the kinds of dihedral angle changes these residues might assume in response to actual protein domain motions associated with binding events. Consequently we propose additional residues from our data that share these time domain profile characteristics, in case they may be potentially associated with phenomena such as allostery or the formation of druggable pockets. We note that dynamic data is particularly appropriate for searching for druggable pockets due to their transient nature [16]. Naturally, future dedicated work would be needed to confirm the significance of our proposed residues, however. It is also worth mentioning that there were no correlations observed between residues identified by previous groups as associated with allostery and drug pocket formation with those residues that had high dihedral angle standard deviations. In all, our present work therefore indicates that there is structured behavior hidden in the time domain dihedral angle fluctuation data of the S protein. We postulate, however, that many of these observations were likely made possible due to the large size of the protein which permitted a wide sampling of frequency components in the dihedral oscillations for both phi and psi. Nevertheless, further studies are needed to confirm if this behavior is universal to all proteins, regardless of size, or if it is unique to the SARS-CoV-2 S protein. We also note that additional studies would be needed to identify whether the DC component in our data is caused by actual physics in the system such as the effects of the quantity of ions and water, temperature, etc… or whether it is purely artifactual (we call to mind how vertical measurements due to gravity in accelerometers such as those found in smartphones can cause large DC components in those systems as well).

